# Towards a unifying phylogenomic framework for tailed phages

**DOI:** 10.1101/2024.10.21.619452

**Authors:** Alaina R. Weinheimer, Anh D. Ha, Frank O. Aylward

**Author notes:** correspondence to (FOA). Author emails: ARW –, ADH –, FOA –.

## Abstract

**Background:** Classifying viruses systematically has remained a key challenge of virology due to the absence of universal genes and vast genetic diversity of viruses. In particular, the most dominant and diverse group of viruses, the tailed double-stranded DNA viruses of prokaryotes belonging to the class Caudoviricetes, lack sufficient homology in the genetic machinery that unifies them to reconstruct inclusive, stable phylogenies of these genes. While previous approaches to organize tailed phage diversity have managed to distinguish various taxonomic levels, these methods are limited in scalability and reproducibility, and they do not include modes of evolution, like gene gains and losses.

**Results:** Here, we present a novel, comprehensive and reproducible framework for examining evolutionary relationships of tailed phages. In this framework, we compare phage genomes based on presences and absences of a fixed set of gene families which is used as binary trait data that is input into maximum likelihood models, which include heterogeneous rates of trait losses and gains. Our resulting phylogeny stably recovers known taxonomic families of tailed phages, with and without the inclusion of metagenomic phages. We also quantify the mosaicism of replication and structural genes among known families. Our results suggest that these exchanges likely underpin the emergence of new families. Additionally, we apply this framework to large phages (>100 kilobases) to map emergences of traits associated with genome expansion.

**Conclusion:** Taken together, this evolutionary framework for charting and organizing tailed phage diversity improves the systemization of phage taxonomy, which can unify phage studies and advance our understanding of their evolution.

## Introduction

Viruses of prokaryotes, often referred to as phages, are ubiquitous and abundant constituents of habitats across the globe [1]. Understanding their evolution and diversity is crucial to characterize their impact on microbial evolution and diversity, which ultimately influences biogeochemical cycles and organismal health that microbes modulate [1,2]. The massive increase of sequencing data and application of metagenomics to a wide variety of environments has vastly expanded our knowledge of viral diversity, and created challenges for classifying uncultivated viruses based on genomes alone. At present, most known phages are tailed, double stranded DNA viruses belonging to the class *Caudoviricetes* (realm Duplodnaviria) and are defined by a small set of hallmark proteins, including a HK97-fold major capsid protein, terminase, portal protein, and prohead protease [3]. The vast diversity of phages within the Caudoviricetes often hinders efforts to assess the phylogenetic relationships between different divergent lineages within this group [3]. This is due to both the highly divergent nature of hallmark proteins within the Caudoviricetes, as well as the highly mosaic nature of phage genomes, in which entire functional modules are often transferred between groups [4]. As a result, lower levels of the Caudoviricetes like orders and families are typically manually proposed based on specific features, and it has remained challenging to assess broader patterns of phage diversity [5].

In previous decades a variety of approaches have been developed to examine broad evolutionary trends across the Caudoviricetes. The Phage Proteomic Tree was among the first workflows designed to assess the phylogenetic relationship between phages by generating trees based on the overall similarity of encoded proteins [6]. This was an important development that opened the door to subsequent genome-based taxonomy approaches. More recently, a variety of methods such as VipTree [7] and VICTOR [8] have adapted similar approaches to larger genomic datasets to examine a much broader range of phages and further streamline taxonomic demarcation of phage lineages. Phylogenetic analysis of sparse multi-sequence alignments has also been proposed as a method to deal with the lack of universal and alignable marker genes in phages [9], albeit based only on reference phage genomes and do not represent uncultivated, environmentally-derived genomes. Lastly, network-based approaches that quantify patterns of gene sharing between genomes and metagenome-derived contigs have also been developed to examine phage diversity to deal with the limited homology of gene sequences among phages [10,11]. While such is informative for addressing the rampant horizontal gene transfer, genetic mosaicism, and incompleteness of uncultivated genomes, many of these network-based methods rely on *de novo* protein cluster or sequence motif generation, which can limit reproducibility of the results as new genomes are input. Moreover, given the mosaicism of phage genomes, it can be difficult to interpret network-based representations of phage similarity.

In this study, we sought to examine the phylogenetic relationships of phages through evolutionary reconstruction of protein family occurrence profiles and to develop a novel approach that could be used for taxonomic delineation of phage lineages. This approach leverages maximum-likelihood phylogenetic reconstructions of Viral Orthologous Group (VOG; vogdb.org) occurrences, thereby preventing the loss of information that takes place when whole-genome distance metrics are used. We identify and benchmark a set of VOGs that consistently recapitulate current taxonomic demarcations within the *Caudoviricetes* and successfully groups together families with known evolutionary links, such as the T4-like supergroup (Straboviridae, Ackermannviridae, and Kyanoviridae) [5,12]. We also apply this method to a set of over 9,000 complete *Caudoviricetes* genomes of diverse metagenomic datasets to provide insights into global tailed phage diversity and phylogenetic structure. Because we use a fixed set of VOGs, our results are reproducible and enable detailed examination of mosaicism across tailed phages that can be used to demarcate broad taxonomic ranks in the future. Our results provide a novel perspective on the evolution of the *Caudoviricetes* and an additional tool for examining evolutionary relationships within this group.

## Results and Discussion

### Evolutionary framework for charting *Caudoviricetes* diversity reveals phylogenetically meaningful protein families

To both maximize and balance diversity for phylogenetic reconstruction, we compiled a representative set of both reference and metagenomic complete genomes of *Caudovircetes* (See Methods). This curation resulted in 2,534 representative, complete *Caudoviricetes* genomes from a total of 19,941 genomes in the INPHARED database (herein called inCaudo genomes downloaded on November 11, 2022; Supplemental Dataset S1), and 9,144 representative, complete *Caudoviricetes* genomes from metagenomes of a variety of environments (herein called metaCaudo genomes) (Supplemental Dataset S1). This resulted in a final database of 11,677 high-quality *Caudoviricetes* genomes that we used for subsequent analyses. To identify a subset of VOGs suitable for phylogenetic inference, we first determined the prevalence of each VOG across our representative set of *Caudoviricetes* genomes. We demarcated four sets of VOGs based on their distribution across genomes: those found in at least 0.25% (n=4,042 VOGs), 0.5% (n=2,114), 1% (n=1,032), and 2% (n=472) of the genomes (Supplemental Dataset S2).

We then ran a phylogenetic reconstruction of only the inCaudo genomes, by generating a binary fasta file based on the presence and absence VOGs from each set and using maximum-likelihood reconstruction that employed a generalized time-reversible model (GTR2-CAT; Figure 1a; see Methods). We found that both the VOG subsets that included VOGs present in 0.25% and 0.5% of *Caudoviricetes* genomes recovered monophyletic clades that corresponded to ICTV-demarcated families (Supplemental Figure S1). Because the 0.5% set produced a tree of higher quality, albeit only slightly (Supplemental Table S1), we proceeded with the 0.5%, also due to its smaller number of VOGs, which would allow better scalability for further analysis. We heretofore refer to this set as the *Caudoviricetes* phylogenetic VOGs (cpVOGs).

**Figure 1.**
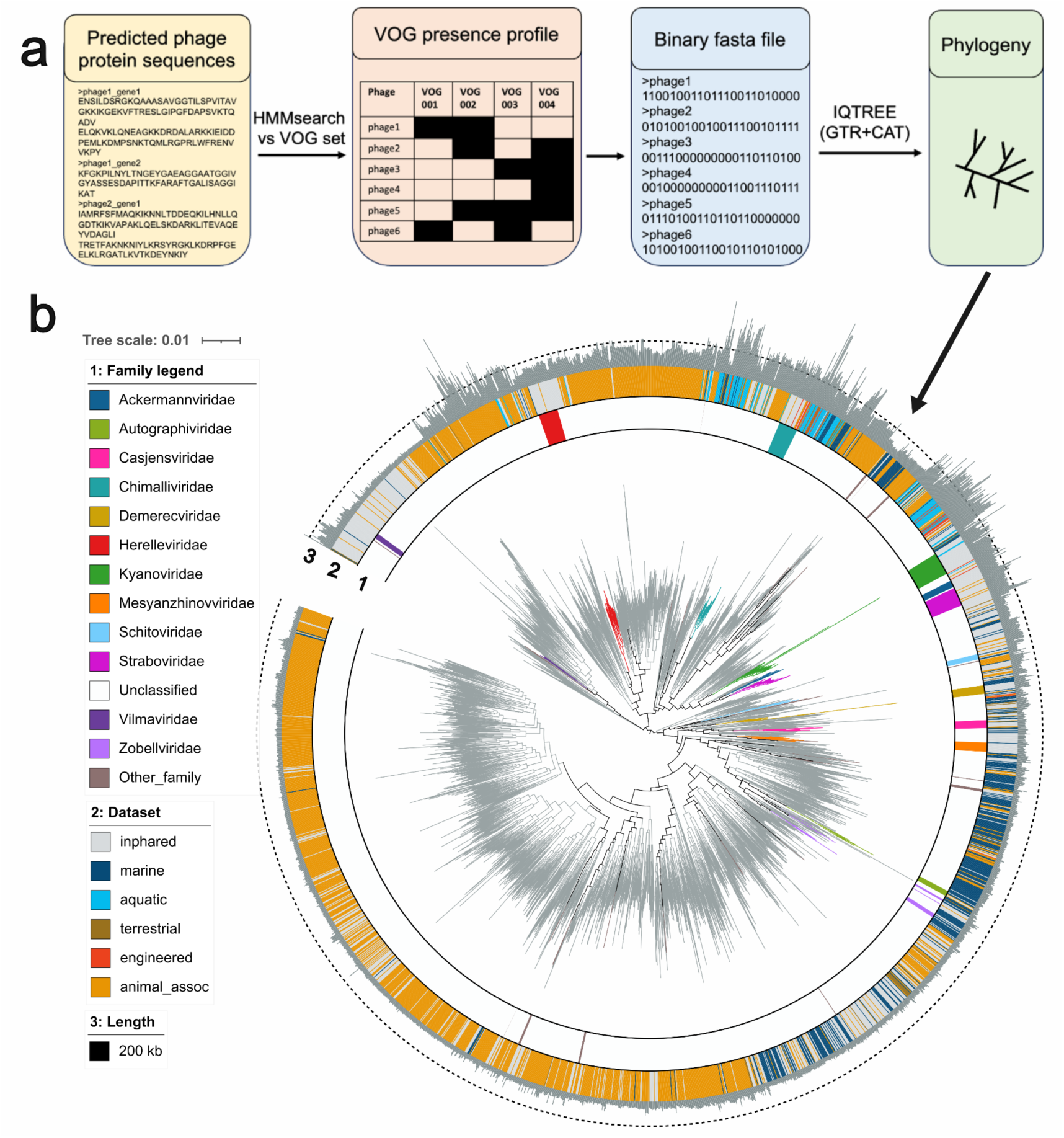
Reconstruction of Caudoviricetes evolutionary history via VOG distribution. (**a**) Bioinformatic workflow used to generate phylogenies from VOG profiles. (**b**) Phylogeny of representative Caudo genomes inferred from their cpVOG profiles generated via the workflow in (**a**). Branches are colored by family, ring 1 color strip is colored by family, ring 2 color strip is colored by source dataset or environmental category, ring 3 indicates genome length with the dashed line corresponding to the jumbo phage (200 kilobase) mark.

To ensure these cpVOGs maintained phylogenetic consistency when metagenome-derived genomes were included, we reconstructed the phylogeny with both the metaCaudo and inCaudo genomes using their cpVOG profiles, again filtering for genomes with at least 5 cpVOGs present (11,646 Caudo genomes). This tree also recovered the monophyly of known families and orders (Supplemental Figure S2). Furthermore, because overrepresentation of different taxa can lead to inaccurate branching [13], we clustered these metaCaudo and inCaudo genomes together based on euclidean distances of cpVOG profiles (see Methods), which resulted in a reduced set of 3,052 representative Caudo genomes (Supplemental Dataset S1). With the binary cpVOG profiles of these representative Caudo genomes, we reconstructed their phylogeny with the same approach as above. This representative tree maintained the recovery of known monophyletic families, further validating our approach (Figure 1b). We compiled this workflow into a python-based tool called VirTree that is available on GitHub https://github.com/faylward/virtree). On the GitHub page, we have also made the cpVOG databases available for download and provided instructions for applying this workflow to a custom set of phage genomes.

### Functions of prevalent cpVOGs suggest widespread temperate lifestyle and reveal uncharted diversity within the Duplodnaviria

We then sought to examine the distribution of cpVOGs across the inCaudo and metaCaudo phylogenies. The class *Caudoviricetes* belongs to the realm Duplodnaviria, or HK97-MCP supermodule, which has distinctive functional genes: HK97-MCP, terminase large subunit (TerL), terminase small subunit (TerS), and portal proteins [3]. While these proteins are relatively conserved, their underlying amino acid sequences are quite divergent, such that no singular orthologous gene group has been found as conserved across the entire *Caudoviricetes* when their genes are clustered together [3]. Likewise, no single VOG was found in every Caudo genome of this study, with the most prevalent VOG found in only 34.7% of genomes (a TerL, VOG00023). This lack of conservation of VOG families across genomes underscores the broad diversity in the *Caudoviricetes*, which continues to challenge the detection and characterization of their diversity in nature and limit the use of single-gene trees for accurately representing their evolutionary history.

Among the most common cpVOG families in the Caudo genomes apart from the TerL VOG00023, were an integrase (VOG 20, 30.2% of genomes) and repressor protein cI (VOG01128 in 24.6%). These genes are used in temperate phages to integrate into the genome of their host (integrase) and prevent transcription of their genes while in the latent state (repressor protein cI) [14]. Notably, these temperate-associated genes were slightly less present in the inCaudo genomes (VOG00020 integrase: 27.6%; VOG01128 repressor protein cI: 19.2%) compared to the metaCaudo genomes (VOG00020: 30.8%, 26.1%), suggesting phages in nature may use a temperate lifestyle more commonly than isolated phages. Furthermore, the presence of the integrase and repressor cI were scattered across the phylogeny (Supplemental Figure S3), suggesting that lysogeny is a highly plastic trait that has evolved in many distinct lineages through acquisition of the necessary machinery for host genome integration.

### Distribution of replication and structural cpVOGs across different families highlights deep mosaicism within the *Caudoviricetes*

Next, we sought to examine the relationship between the evolutionary patterns revealed through marker gene phylogenies and our method of phylogenetic inference based on VOG profiles. For this we performed phylogenetic inference on four broadly-represented VOGs that are found across a range of *Caudoviricetes* for which ICTV-demarcated families have been constructed. We examined trees made from the TerL (VOG00012) and family B polymerase (VOG00800) found in the T4 supergroup families of Ackermanviridae, Kyanoviridae, and Straboviridae, and we compared these to trees made with cpVOG-based inference of the same genomes (Figure 2ab). Similarly, we also compared the TerL (VOG000141) and family A DNA polymerase (VOG00013) found in a wide range of T7-like phage families and their relatives and made the same comparison to cognate cpVOG trees (Figure 2cd). In all comparisons, we generally found that both marker gene and cpVOG trees recovered monophyletic families. The main exception was the family A polymerase tree of the Autographiviridae, which recovered two distinct and divergent groups, while the cpVOG tree recovered a single Autographiviridae family. Detailed inspection of the VOG00013 phylogeny confirmed that different lineages within the Autographiviridae indeed encode polymerases that belong to two distinct clades (Supplemental Figure S4). Moreover, individual marker genes sometimes provide conflicting results; for example, the TerL tree of T4-like phages suggested that the Ackermannviridae are basal to the Straboviridae and Kyanoviridae, but the cpVOG trees and family B polymerase trees suggest a closer evolutionary link between the Ackermannviridae and Kyanoviridae. Collectively, these results confirm that individual marker gene trees can provide conflicting results, and the cpVOG-based phylogenetic reconstruction that we present here may offer a useful alternative.

**Figure 2.**
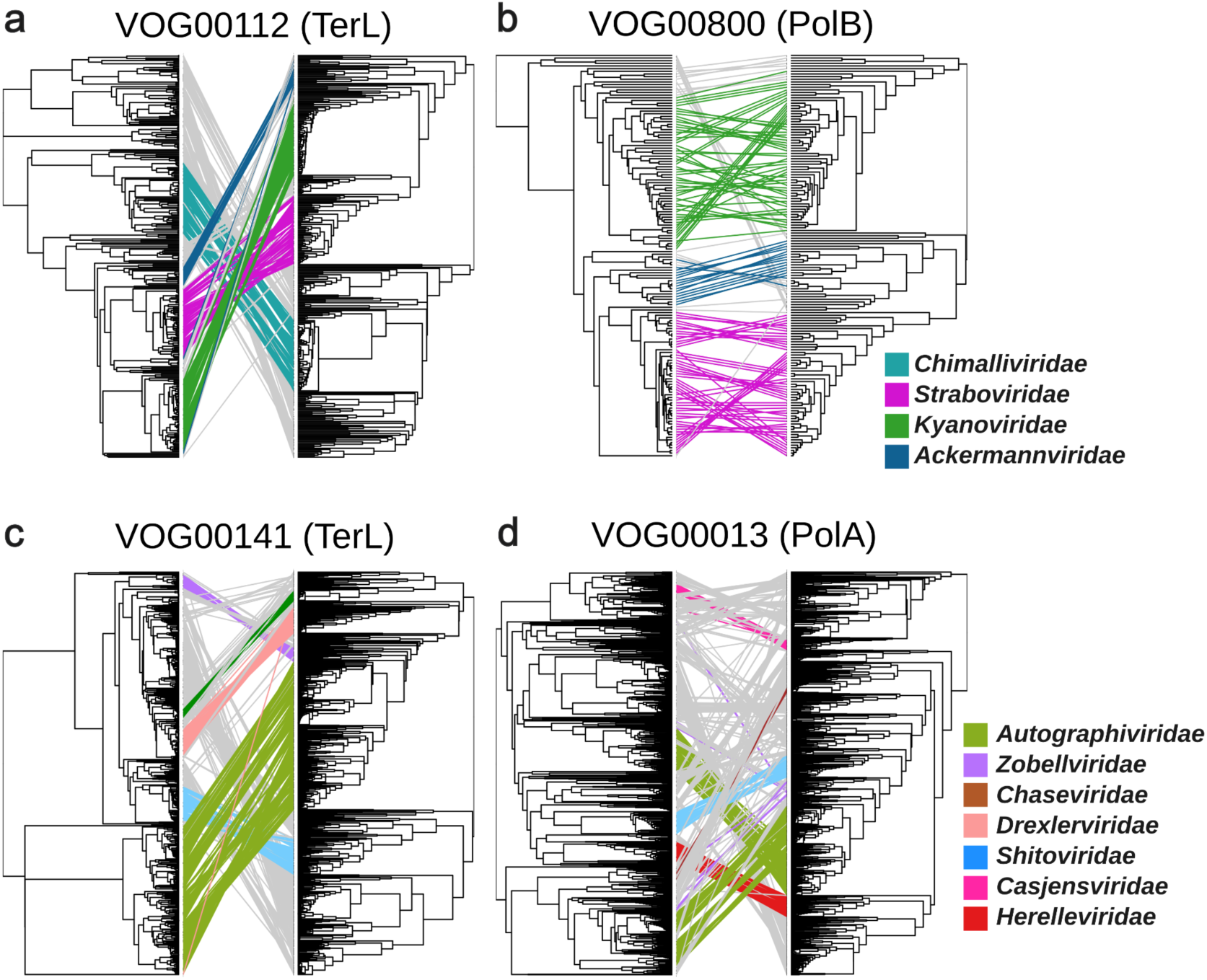
Tanglegram comparison of single gene trees with cpVOG-inferred phylogenies. Left side phylogeny corresponds to the phylogeny inferred from the alignment of amino acid sequences of a given VOG encoded by the inCaudo genomes. Right side corresponds to the phylogeny inferred from the cpVOG profiles of inCaudo genomes encoding the given VOG. Branches are colored by family. (**a**) Tanglegram of the TerL VOG00112 (**b**) Tanglegram of DNA polymerase B VOG00800 (**c**) Tanglegram of TerL VOG00141 (**d**) Tanglegram of DNA polymerase A VOG00013.

Our phylogenetic reconstruction of cpVOG profiles demonstrates that this approach robustly recapitulates ICTV-approved family demarcations, and we therefore examined the distribution of key protein families in these lineages (Figure 3a). The Ackermanviridae, Kyanoviridae, and Straboviridae families all fall within the informal “T4 supergroup” that has been described previously [5], and, consistent with, this they form a monophyletic lineage in our cpVOG tree. These families also encode the same family of TerL, MCP, and family B polymerase (VOG00112, VOG00292, and VOG00800, respectively; Figure 3a), underscoring their clear evolutionary affinity. Interestingly, the *Chimalliviridae*, a recently demarcated family of “jumbo” phages (genomes over 200 kilobases [15]) that have particularly large genomes [5,16], encode the same family of TerL but distinct MCP and polymerase VOGs, indicating that this family has a distant evolutionary affinity for the T4 supergroup. The Zobellviridae, Autographiviridae, and Shitoviridae also have the same families of TerL, MCP, and polymerase (family A), indicating that these lineages also share an ancient evolutionary link, while families such as the Chaseviridae or Drexlerviridae encode only some of these.

**Figure 3.**
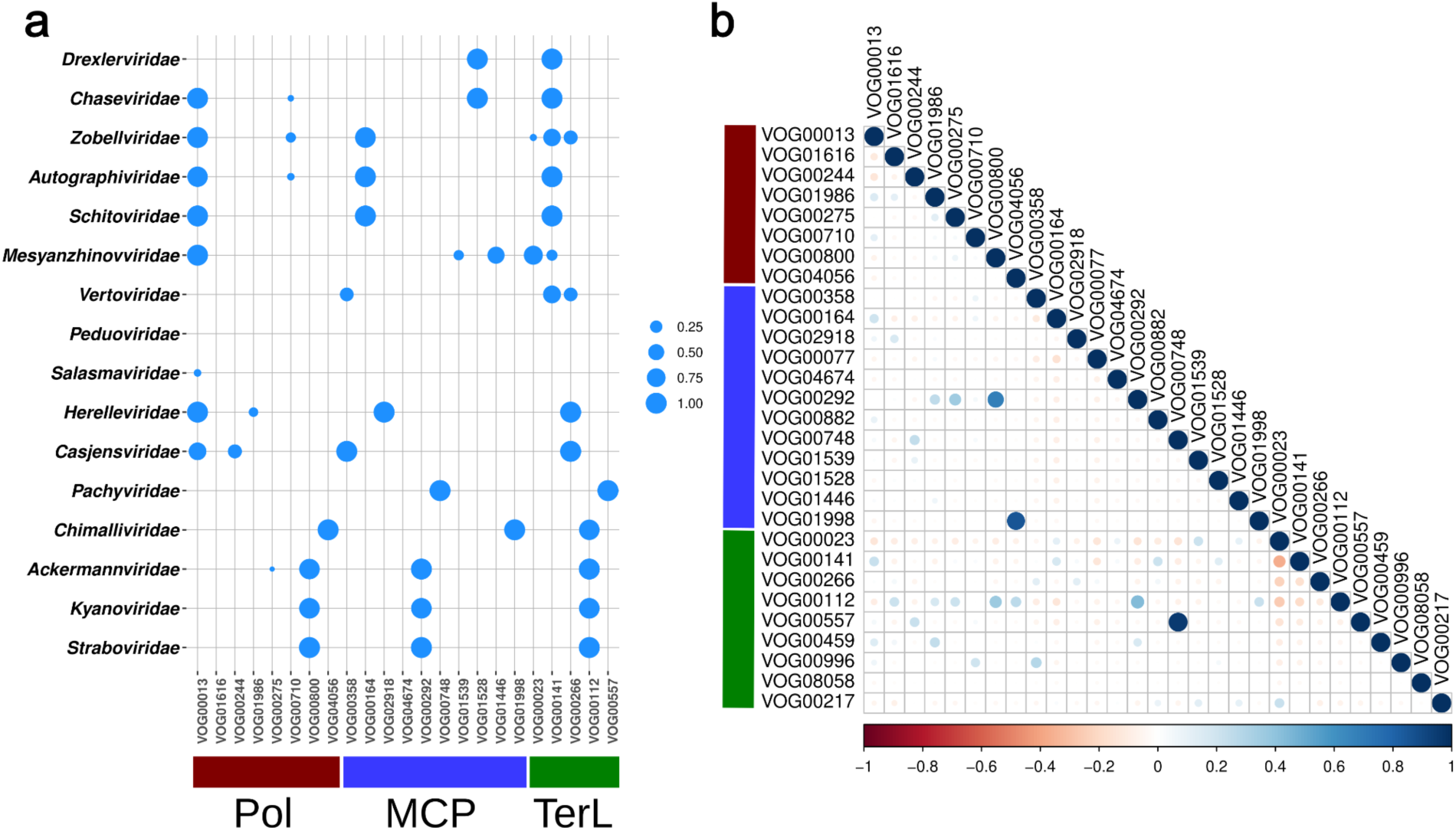
Distribution of VOGs belonging to functional groups (Pol, MCP, TerL) across families. (**a**) Bubbleplot in which the size of each point corresponds to proportion of genomes in the taxonomic family that encode a VOG in the different functional groups. **(b**) Correlogram depicting the co-occurence of different VOGs with each other of the different functional groups. Red points indicate a negative correlation between the occurrence of two VOGs, white indicates no correlation, blue indicates a positive correlation. Color intensity corresponds to the strength of the correlation. Abbreviations: Pol - DNA polymerase, MCP - major capsid protein, TerL - terminase large subunit.

We also examined patterns of co-occurrence in cpVOGs to assess if this could reveal trends in deep phage evolution (Figure 3b). In some cases these results are straightforward to interpret. For example, the TerL VOG00112 and MCP VOG00292 are found in the same families in the T4 supergroup and underpin the morphogenetic module in those families. Other examples are less clear, however. For example, the family A DNA polymerase VOG00013 co-occurs with the MCP VOG00141. Although these protein families are involved in distinct functions (genome replication vs. virion structure), their co-occurrence suggests deep evolutionary links between many disparate lineages of phages that use a family A DNA polymerase.

To examine the deep roots of mosaicism in *Caudoviricetes,* we examined the frequency at which different families of TerL, MCP, and DNA polymerases cpVOGs occurred in the same genome of both cultivated and uncultivated phages (Figure 4). These patterns show that these marker genes have been exchanged extensively throughout the diversification of the Caudoviricetes, leading to a widespread mixing of protein family combinations. This is consistent with the extensive level of deep genomic mosaicism that has long been recognized in tailed phages [4,17], and it suggests the recombination of morphogenetic versus informational modules is a mechanism through which new phage families evolve [17]. Viewed in this way, the class Caudoviricetes can be described as a “melting pot” in which replicative and morphogenetic modules have been extensively swapped throughout the emergence of the major groups. Larger phages, however, may experience more evolutionary constraints in this regard because they are “locked in” to specific replisomes and capsids that are suited for their large genomes [18,19]. By contrast, distinct lineages of phage in the 20-80 kb genome size range, may be able to swap capsids or polymerases with relative ease. This may partly explain the widespread mosaicism observed in smaller phages compared to the relatively consistent structure seen in larger phages (i.e. the T4 supergroup) [18,19].

**Figure 4.**
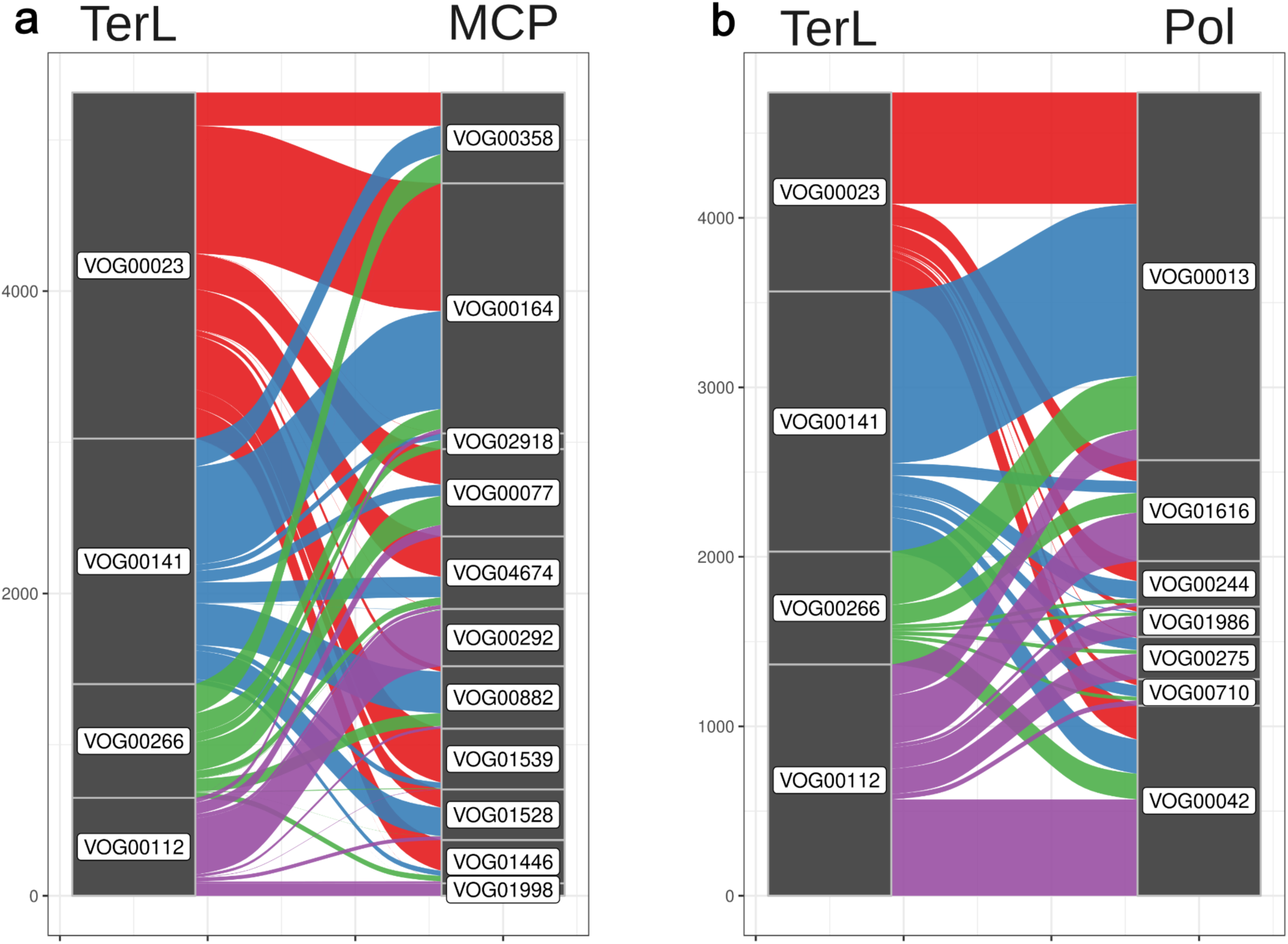
Alluvial plots comparing exchange and co-occurence of VOG families of different functional groups within Caudo genomes. where the left stratum corresponds to VOGs belonging to one functional group and the right to another. Lines are drawn based on their co-occurrence in a Caudo genome. Lines are colored by left stratum’s VOG. (a) contains TerL VOG families on the left and MCP VOG families on the right stratum. (b) contains the same TerL families from (a) on the left and DNA polymerase families on the right stratum.

To quantify the extent of mosaicism and conserved genome structure in phages across different genome size ranges, we identified sets of VOGs that co-occur in the same genomes (Pearson’s Correlation R > 0.7 in our Caudo set of 11,677 genomes). We identified 57 sets of between 3 to 60 VOGs that represent components of consistent genomic architecture in our dataset (Supplemental Dataset S2). Importantly, 67.7% of phage genomes > 100 kbp in length contained at least one of these conserved VOG sets despite comprising 13.8% of the Caudo genomes examined, compared to just 31.6% for phage genomes < 100 kbp in length while comprising 86.2% of Caudo genomes examined. Thus, larger phage genomes therefore contain more conserved genomic structure that can be readily detected in VOG profiles. Moreover, we found that overall VOG profile correlation distributions were higher for larger phage genomes (Figure 5), which we would expect if there were more conserved elements of genome composition across these phages. Together, these findings corroborate the relatively more stable genome architecture of larger phages compared to smaller ones. This is consistent with studies of T4-like phages that identified both a conserved genomic backbone as well as highly plastic variable regions [12]. Moreover, virulent phages have been described as “low gene flux” compared to temperate phages, but this pattern may be driven in part by the typically larger genomes of virulent phages [18,19]. We suggest that larger phage genomes tend to contain conserved architecture simply because larger genomes are not as amenable to large-scale mosaicism. In other words, larger phages may be “locked in” to a particular genome structure, potentially due to more complex protein-protein interactions that take place during their infection cycles. Meanwhile, smaller phages may more readily recombine morphogenetic and replication modules between lineages, leading to novel genome architectures.

**Figure 5.**
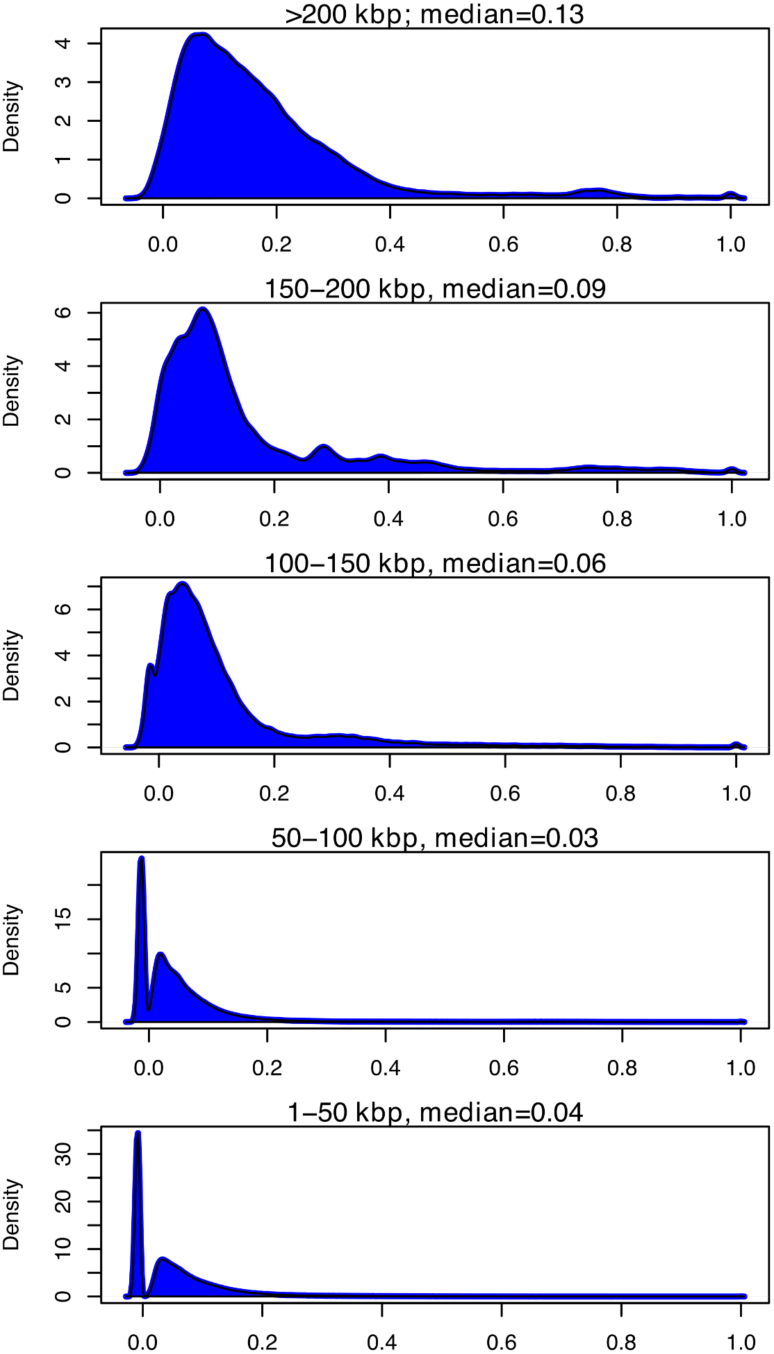
Density of correlations between VOG profiles of genomes within different genome length groups. The x-axis is the correlation between VOG profiles of genomes within that group. The y-axis is the frequency of that correlation within the group calculated with the density function in R. The titles correspond to the genome length group in kilobases (kbp) and the median is the median Pearson correlation of that group.

### Case study on large phages reveals multiple independent origins of jumbo phages

Finally, we used this cpVOG-based approach to examine the evolution of genome gigantism across the *Caudoviricetes* by performing an in-depth analysis on phages with genomes of at least 100 kilobases. A previous study on the evolution of large phages using dendrograms of gene-sharing distances within cultured phages posited that large genomes evolved from smaller relatives in multiple, independent occasions [20]. Likewise, our cpVOG phylogenetic approach estimates at least 19 emergences of jumbo phage clades, highlighting their multiple, independent origins from smaller phages (Figure 6). Our tree puts these emergences into broader context with the inclusion of environmental phages.

**Figure 6.**
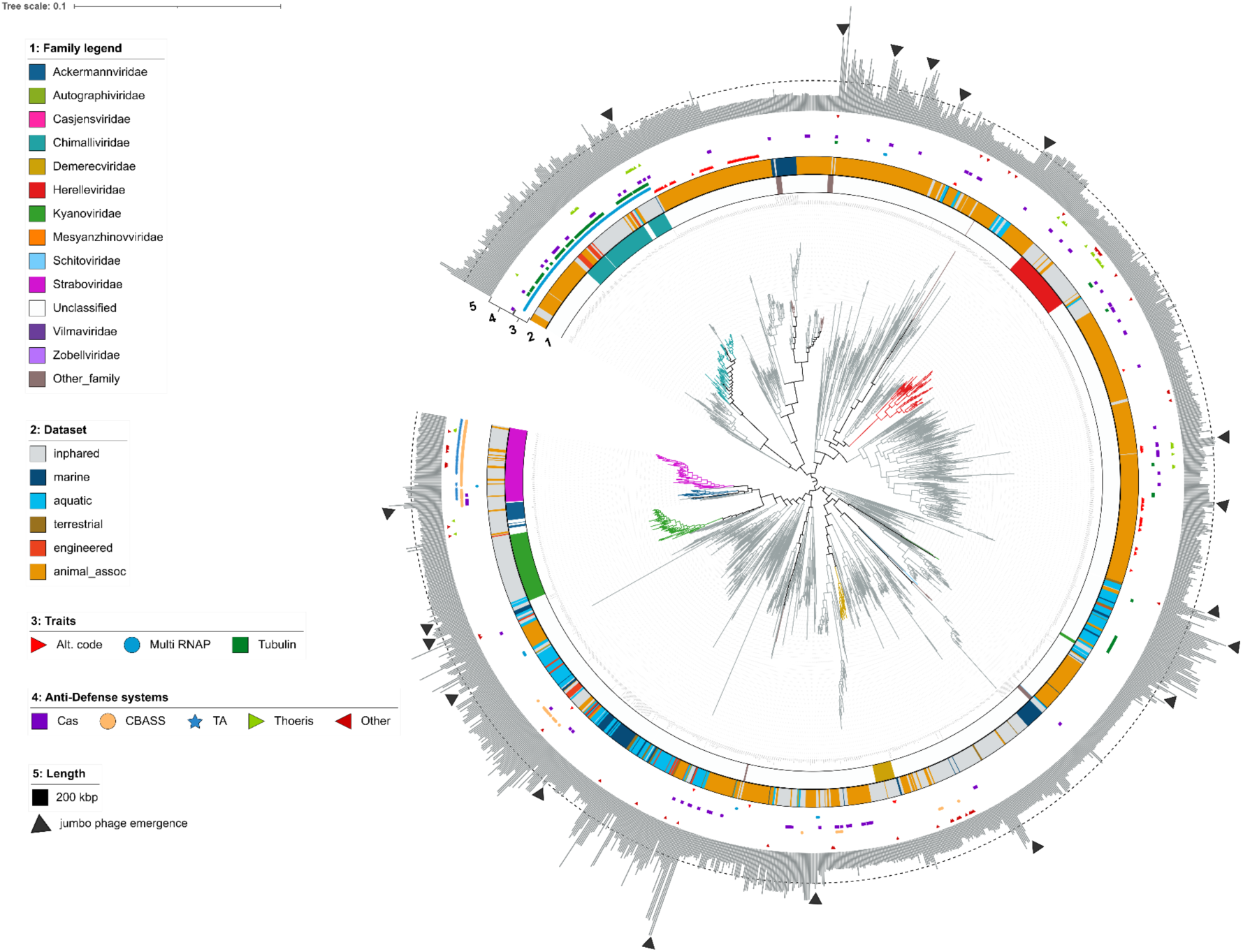
Phylogeny of large (> 100 kilobases) Caudoviricetes genomes. Branches are colored by family. Ring 1 corresponds to family; ring 2 corresponds to environment; ring 3 corresponds to the presence of the following features: alternative genetic code (Alt. code), multisubunit DNA-dependent RNA polymerase (Multi RNAP), and tubuiln; ring 4 corresponds to the presence of different APIS genes (Cas - anti-CRISPR Cas, TA - toxin-antitoxin); ring 5 displays genome length in kilobases (kbp) with a dashed line corresponding to 200 kilobase and black triangles marking emergences of jumbo phages (> 200 kb).

Several recent studies have identified features such as alternative genetic codes, multisubunit DNA-dependent RNA polymerases (multi-RNAP), and tubulin that are associated with lineage of large phages [16,21,22], and we therefore sought to evaluate the prevalence of these traits in our large phage tree (Figure 6; Supplemental Dataset S3). This analysis revealed that tubulin homologs appear to be a distinguishing function of the *Chimalliviridae* [23], which are known to use this gene as part of their “phage nucleus” structure that they form during infection, although a handful of large phages scattered across the tree encode it as well. Multi-subunit RNAP was also prevalent among the *Chimalliviridae* and the *Straboviridae*. Although these families share the same family of TerL (VOG00112), they are otherwise quite divergent and are placed in distinct areas of our VOG profile-based trees, consistent with the view that these lineages may have convergently acquired multi-subunit RNAP. Finally, the use of an alternative genetic code has been a feature uncovered in very large phages (over 500 kilobases) called Lak phages [22]. This strategy is hypothesized to facilitate phage replication [24]. We examined the distribution of large phages using alternative genetic codes across our tree and found such only among two clades of metagenome-assembled genomes, which contained those under 200 kilobases, as well as those above (Figure 6). The two clades were separated by several established families, suggesting that the use of alternative genetic codes among large phages has emerged on independent occasions.

Lastly, several studies have noted that large phages appear to encode a wide array of anti-defense systems that obfuscate host defense systems [25,26]. We therefore examined the distribution of known anti-defense systems across large phages, including anti-CRISPRs, CBASS, toxin-antitoxin (TA), Theoris, and others (see Methods; Supplemental Dataset S3). Our results found that these anti-defense systems are broadly scattered across our large phage tree with a high degree of variability within clades, suggesting that they are highly mobile in the virosphere and that their distribution may be driven by recent host-virus co-evolution. Importantly, we found that anti-defense systems were statistically enriched in phages with genomes >200 kbp compared to smaller phages (Mann-Whitney U test, p < 0.001), suggesting that the acquisition of these systems may be a driving force behind the evolution of genome gigantism (Supplemental Dataset S3).

## Conclusions

Examining evolutionary relationships between tailed phages has always been difficult owing to the high level of genomic mosaicism and sequence divergence in this group [3,17]. As a result, examining evolutionary relationships deeper than the family level has remained challenging. In this study, we present a novel approach to examine evolutionary relationships of tailed phages (*Caudoviricetes*) based on the phylogenetic reconstruction of protein family occurrences. We show that this approach reproduces ICTV-approved families as well as broader evolutionary relationships that are well-known, such as those that link the T4-like families *Straboviridae, Kyanoviridae*, and *Ackermannviridae*. Moreover, we show that this approach can be used to identify hallmark protein families of phage lineages and identify instances of deep mosaicism across families. The deep mosaicism that is apparent in the *Caudoviricetes* suggests that new phage lineages emerge through recombination of the morphogenetic and replicative modules of disparate lineages, and analysis of these patterns may yield insight into ancient recombination events that have led to the rise of present-day lineages in the virosphere.

Additionally, our framework provides a new, robust method for placing novel phages into a phylogenetically meaningful context of known diversity, as well as identifying functional modules that are found in other lineages, thereby facilitating taxonomic delineation of novel groups. Given that the vast majority of phages currently do not fall within lower-level taxonomic groups, approaches that can compare a large number of genomes regardless of the presence of specific marker genes and provide reproducible results are highly valuable. As known viral diversity continues to expand with increasing environmental sequencing data, defining an objective and scalable approach to charting viral diversity and evolutionary relationships, such as presented here, is necessary for progressing a synoptic understanding of the virosphere.

## Methods

### Dataset compilation and dereplication

Reference *Caudoviricetes* were downloaded from the INPHARED database (November 2022;[27]) based on assignment to a Caudoviricetes family list via the ICTV as of November 2022 (Supplemental Dataset S1), which included 19,941 genomes. These were de-replicated at 90% ANI with dRep (v 3.2.2; options: dereplicate --ignoreGenomeQuality -p 32 -l 5000 -pa 0.90 –SkipSecondary; [28]), which resulted in 3,263 representative genomes. These were then filtered for those encoding at least 20 kilobases, resulting in 3,125 genomes. As some phages on INPHARED are metagenomic, phage genomes that were not designated as complete in their name and/or labeled metagenomic were further filtered out, resulting in 2,534 genomes. Metagenome-derived *Caudoviricetes* genomes over 20 kilobases in length were downloaded from human gut samples [29,30], a jumbo phage study [21], marine samples [31–36], peat samples [37] and two wastewater studies [38,39]. Metagenomic genomes were determined as *Caudoviricetes* either based on the classification specified in the study or by having a maximum score for dsDNAphages using VirSorter2 ([40],v 2.2.3, default options). Completeness of metagenomic genomes were either specified in the study or contained DTR or ITR as detected by CheckV in the “complete_genomes.tsv” output file ([41], checkv end_to_end; default options). Apart from the jumbo phage genomes, to reduce redundancies in the collective dataset for downstream analysis, each dataset with over 100 complete Caudoviricetes genomes over 20 kb in length were de-replicated with dRep using a mash distance of 0.9 (dRep dereplicate –ignoreGenomeQuality -l 20000 -pa 0.9 --SkipSecondary). The 336 Al-Shayeb et al. 2020 jumbo phage genomes were not de-replicated to retain jumbo phage diversity for genome size analyses. After these steps, 9,145 metagenomic genomes resulted, with 3,735 from marine metagenomes, and 4,898 from human gut metagenomes, 164 from peat, and 12 from wastewater. These 9,145 metagenomic phage genomes (metaCaudo) were combined with the 2,534 INPHARED phage genomes (inCaudo) for a total of 11,677 complete Caudoviricetes genomes over 20 kilobases (Supplemental Dataset S1).

### Protein prediction

Some phages have been found to use genetic codes alternative to the Standard 12 [22]. To determine the genetic code of the phages in our dataset, we ran prodigal on all genomes with all codes available in prodigal (1, 2, 3, 4, 5, 6, 9, 10, 11, 12, 13, 14, 15, 16, 21, 22, 23, 24, 25) [42]. If the genetic density of a genome was over 80% using the Standard 11, we did not test alternative codes to reduce the potential over-prediction of short genes. For those under 80%, we predicted genes with the remaining codes in prodigal and selected the prediction with the highest genetic density. The python script to generate the optimal genetic code is deposited in the script “genetic_code_tester.py” in the Caudoviricetes phylogeny dataset repository on Zenodo. Genetic codes of genomes are in Supplemental Dataset S1.

### Generating gene content profiles with the VOG database

We aligned amino acid sequences of each genome to HMM profiles from the Virus Orthologous Groups (VOG) database (vogdb.org, release 214; Supplemental Dataset S2) via hmmsearch (E value < 10^−5^ HMMER v3, hmmer.org, [43]) as it comprehensively and specifically generates gene families of all virus types on RefSeq at the time of construction. We then selected each gene’s best VOG hit based on the highest bitscore and combined these hits of each genome into a matrix of all genomes with all VOGs found in the genomes (11,677 genomes (rows) by 35,106 VOG families (columns); table in Caudoviricetes phylogeny dataset repository on Zenodo). To identify the most phylogenetically meaningful VOGs, we first screened for VOG families present in a minimum percent of Caudo genomes: 2%, 1%, 0.5%, and 0.25% of genomes (Supplemental Dataset S2; script subset_xpercent_vog_prof.py in the Caudoviricetes phylogeny dataset repository on Zenodo), which we refer to as VOG subsets. We then re-annotated the Caudo genomes with each of these VOG subsets via HMM searches to generate the gene profiles of the genomes (Supplemental Datasets S2; profiles in Caudoviricetes phylogeny dataset repository on Zenodo).

### Determining VOG subset for phylogenetic analyses and benchmarking

For phylogenetic reconstruction of the genomes based on each VOG subset, we first filtered the genomes for those that contained at least 5 hits to a given VOG subset. We then used the VOG profiles to generate binary fasta files of the Caudo genomes in which each header was a genome and the characters were 0’s and 1’s representing the presence and absence of different VOGs (Figure 1; see Caudoviricetes phylogeny dataset repository on Zenodo for binary fasta files). To assess the phylogenetic signal of each VOG subset, we first only used binary fasta files of the inCaudo genomes (INPHARED Caudoviricetes genomes). Accordingly, several VOGs of the subsets 1-0.25% were absent in the INPHARED genomes and thus removed from their binary fasta files for these phylogenies. For the 1% VOG subset this was VOG07561 (Iron-sulfur cluster insertion protein ErpA). For the VOG 0.5% subset, an additional VOG was absent (VOG15190 a “tRNA 4-thiouridine(8) synthase”). For 0.25% VOG subset, in addition to those two VOGs, the six following VOGs were absent: VOG00522, VOG12113, VOG20148, VOG23005, VOG34688, VOG34865 (See Caudoviricetes phylogeny dataset repository on Zenodo). A phylogeny of each VOG subset’s binary fasta file of the inCaudo genomes was generated with IQ-TREE (-st BIN -m GTR2+ASC -nt AUTO -bb 1000 -wbt --runs 5; version 2.2.5; [44]). Tree certainty and internode certainty was calculated with RAXML (8.2.12) [45] (raxmlHPC -f i -m GTRCAT) (Supplemental Table S1; Caudoviricetes phylogeny dataset repository on Zenodo for treefiles). Trees were visualized in iTOL [46] and colored according to family designation via INPHARED downloaded in February 2024 (Supplemental Dataset S1) to be examined for monophyly. The VOG subsets that led to monophyls of known families were those found in 0.5% and 0.25% of Caudo genomes (Supplemental Figure S1. The 0.5% was then used for downstream analyses as this VOG subset had slightly higher tree certainty and relative tree certainty values and contained half as many VOGs as the 0.25% VOG subset to enhance computational efficiency of annotation and tree reconstruction (Supplemental Table S1). With the 0.5% VOG subset, the VOG profiles of all inCaudo and metaCaudo genomes with at least 5 different VOGs detected in a genome (11,621 Caudo genomes) were used to generate the binary fasta of VOG presence and absences input into a larger phylogeny using the same IQ-TREE settings as the inCaudo-only tree. Due to the large number of leaves, many of the nodes were collapsed (those containing 200-500 genomes). This phylogeny was colored by family and retained the monophyls (Supplemental Figure S2). Nonetheless, due to the constraints of visualization and potential for the overrepresentation of certain taxa unbalancing the tree [13], we clustered the Caudo genomes based on their VOG profiles of the 0.5% VOG subset. For this clustering, with R (v4.2.2) [47] in RStudio (version 2023.06.2+561) [48], we generated a pairwise distance matrix of the genomes based on the Euclidean distances of their 0.5% VOG subset profiles (base R function dist(method= “euclidean”)). Next, we generated a hierarchical clustering of the genomes based on the distance matrix with the base R function hclust(method = “average”). We then selected branches in the hierarchy with a minimum height of 5 (cuttree(hcut=5)) which resulted in 3,052 representative genomes of each cluster (Supplemental Dataset 1). We then generated a binary fasta file of the representative genomes from their 0.5% VOG subset profile to reconstruct the phylogeny of these genomes with the same IQ-TREE settings before (See Caudoviricetes phylogeny dataset repository for related files). This phylogeny was then visualized in iTOL and colored based on genomic features of family classification, dataset source, and genome length (Figure 1b).

### VOG distribution analyses and tanglegrams

A bubbleplot and correlogram of VOG presence in different inCaudo genome families was visualized in R based on the cpVOG profiles of genomes using ggplot2 [49] in Rstudio. Alluvial plots were generated using the geom_alluvium command in the R package ggalluvial [50]. Tanglegrams were generated to compare the consistency of single gene trees and our VOG profile phylogenies. For the single gene trees, the amino acid sequences with homology to the VOG of interest within inCaudo genomes were compiled and aligned with Muscle5 [51]. Alignments were trimmed for regions that contained >90% gaps with trimAl (-gt 0.1; v1.4.rev15; [52]). Phylogenies of the trimmed amino acid sequence alignments were generated with IQ-TREE (-bb 1000 -T AUTO -wbt -m LG+F+R10 --runs 5). Phylogenies of the VOG binary profiles of the corresponding inCaudo phages containing the VOG of interest were generated from the binary fasta file of these profiles using IQ-TREE (-alrt 1000 --fast -T AUTO -st BIN -m GTR2+ASC -wbt --runs 5). The tanglegrams were visualized and the degree of entanglement between trees was calculated in R with the tanglegram function in the package dendextend [53]. All data related to the tanglegram analyses are in the *Caudoviricetes* phylogeny datasets repository on Zenodo.

For the whole-genome correlation analysis, we calculated pairwise Pearson correlation coefficients from the VOG profiles of each genome in the Caudo dataset (11,677 phages). We then generated density plots of the correlation distributions using the density() function in base R, and then plotted these for different phage genome size categories (see Figure 5). To identify conserved sets of VOGs that co-occur, we calculated pairwise Pearson correlation coefficients for VOGs from the same matrix as described above, and then used the hclust function in base R with the average linkage agglomeration method to assess VOG co-occurrence. We inspected the hclust dendrogram and cut it at a value of 0.7 to recover clusters of VOGs that tend to co-occur across phage genomes, which resulted in 57 sets of three or more VOGs. To calculate the number of phages in which these VOG sets occur, we only considered genomes for which >80% of the VOGs were present (Supplemental Dataset S2).

### Examining large phage genomes

All phage genomes larger than 100 kilobases in length were included (Supplemental Dataset S3). A phylogenetic tree was calculated with IQ-TREE based on the binary VOG profile of the 0.5% set (-st BIN -m GTR2+ASC -T AUTO -bb 1000 -wbt --runs 5). The resulting tree was visualized in iTOL. To identify the presence of tubulin protein and multi-subunit RNA polymerase, we searched phage proteins against custom HMM profiles (on data repository) using HMM searches (E value < 10^−5^, bitscore > 30 (hmmsearch) (Supplemental Dataset S3)). Anti-prokaryotic immune system (APIS) proteins were detected in the large phages with HMM searches (HMMERv3, hmmer.org) and DIAMOND searches [54] (E values < 10^−5^) on phage proteins against the dbAPIS [25], a database of 92 APIS families and 90 anti-CRISPR protein families. Best hits were determined by bitscore. We reported only hits with consistent matches between the two searches (Supplemental Dataset S3).

## Abbreviations

VOG: Virus Orthologous Group
Caudo: all complete Caudoviricetes genomes used in this study
inCaudo: INPHARED Caudo genomes
metaCaudo: metagenome-assembled Caudo genomes
cpVOG: Caudoviricetes phylogenetic VOG
HMM: Hidden Markov Model
APIS: anti-prokaryotic immune system

## Availability of data and materials

Supplemental Figures S1-S4, Supplemental Table S1, and Supplemental Dataset legends are in the Supplemental Information file. All datasets generated and/or analyzed during the current study are available in “Caudoviricetes phylogeny datasets” repository on Zenodo here: https://shorturl.at/JvYBb. Code used in this manuscript is in the VirTree repository on Github here: https://github.com/faylward/virtree.

## Acknowledgements

We thank members of the Aylward Lab for helpful feedback and Evelien M Adriaenssens for thoughtful discussions on phage evolution and the T4 supergroup. This work was performed using compute nodes available at the Virginia Tech Advanced Research and Computing Center.

## Author contributions

FOA and ARW conceived and designed the study. ARW compiled the genomes, benchmarked the workflow, constructed figures, and performed analyses. FOA performed analyses and constructed figures. ADH performed analyses and constructed figures related to the large phage genomes and family designations. All authors drafted and revised the manuscript.

## Consent for publication

Not applicable

## Ethics approval and consent to participate

Not applicable

