## Supplemental Information for "Towards a unifying phylogenomic framework for tailed phages"

**Contents:**

**Supplemental Dataset Legends:**

**Supplemental Dataset 1:** genome dataset information (download location/dates, publication, number of min 20kb, number of complete, number of Caudoviricetes); genome metadata (env, dataset, length, etc.); membership in clustering analysis

**Supplemental Dataset 2**: VOG v214 version information, with accessions' descriptions, species counts, etc. downloaded from VOGDB.org. Table containing which VOGs are in which subset of genomes (0.25%, 0.5%, 1%, 2%) and the tree quality information resulting from each subset.

**Supplemental Dataset 3**: Large phage ( > 100 kilobases) information; table with genomes and their length, presence of traits in Figure 6 ring 3; anti-prokaryotic immune systems genes detected with DIAMOND and HMMsearches; and presence in large genomes visualized in phylogeny Figure 6 ring 4.

**Data repository:** Zenodo with binary fasta files, tree files, amino acid files for tanglegrams, and other related files. Link: [https://shorturl.at/JvYBb](https://shorturl.at/JvYBb)

**Supplemental Figures S1-S4.**

**Supplemental Table 1**

**Supplemental Figures:**

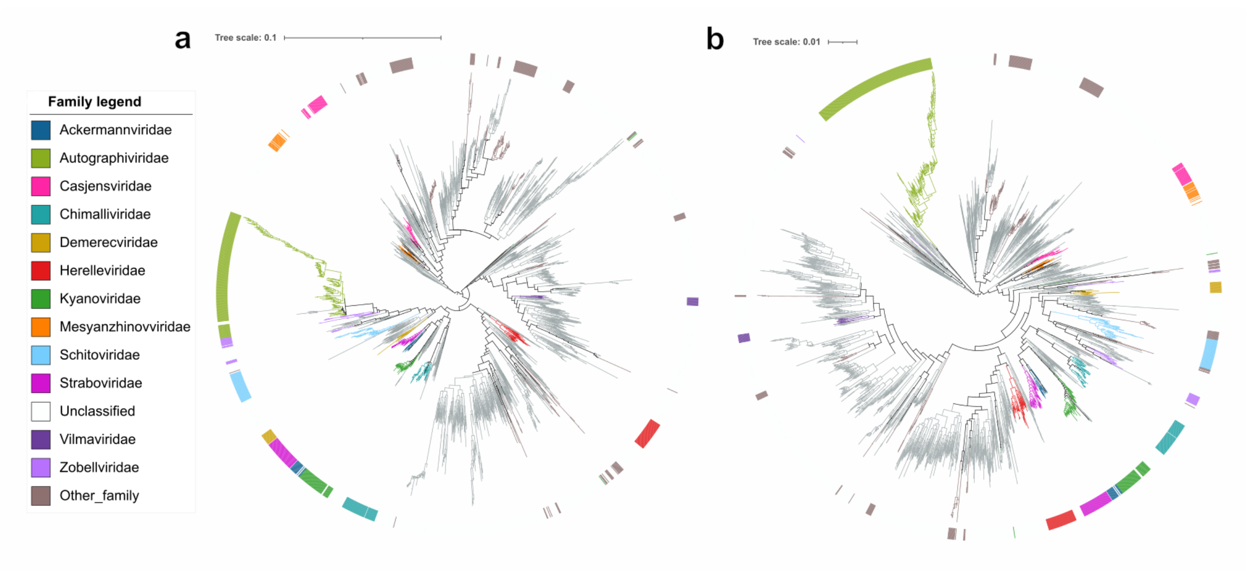

**Supplemental Figure S1.** Phylogenies of INPHARED genomes with at least 5 hits to the 0.5% VOG subset (**a**) or 0.25% VOG subset **(b**). Branch colors and outer ring color correspond to ICTV family. Trees are both rooted at the midpoint.

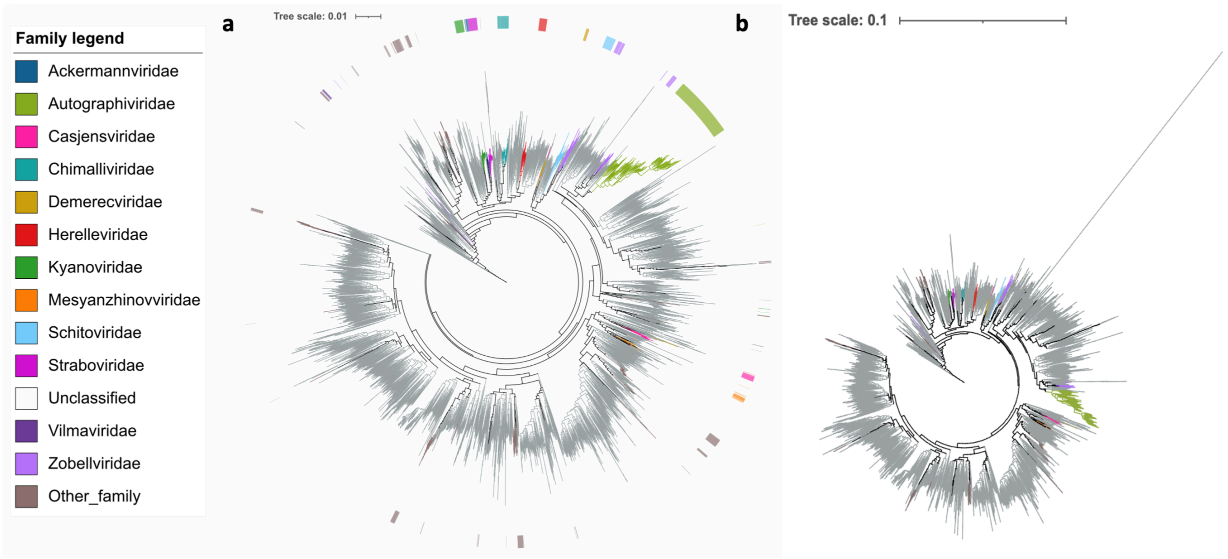


**Supplemental Figure S2**. Phylogeny of all Caudo genomes examined that contained at least 5 hits to the 0.5% VOG subset (11,621 Caudo genomes). Branch colors and outer ring color correspond to ICTV family. Nodes with 200-500 members were collapsed for visualization. (**a**) phylogeny with genome QGNH01001383.1 of the peat metagenomic dataset excluded as its branch length was exceedingly long and shrunk the visibility of the other leaves and their family designations. (**b**) phylogeny with QGNH01001383.1 included.

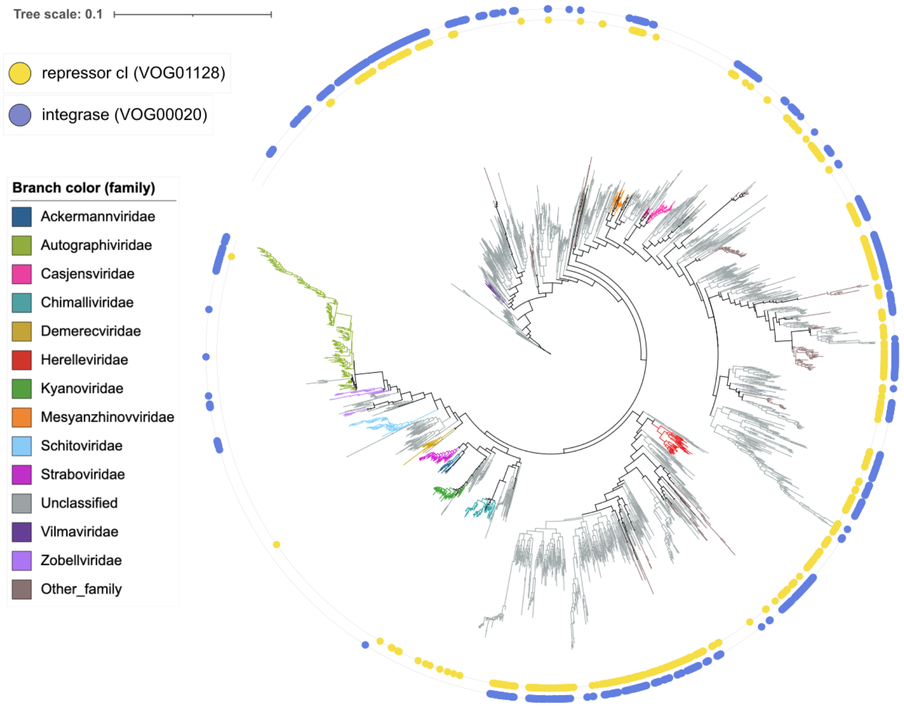


**Supplemental Figure S3.** Representative Caudo 0.5% VOG subset phylogeny (3,052 genomes), with branches colored by family, yellow dots in outer ring corresponding to the presence of the repressor cI VOG (VOG01128) and purple dots to the presence of the integrase VOG (VOG00020).

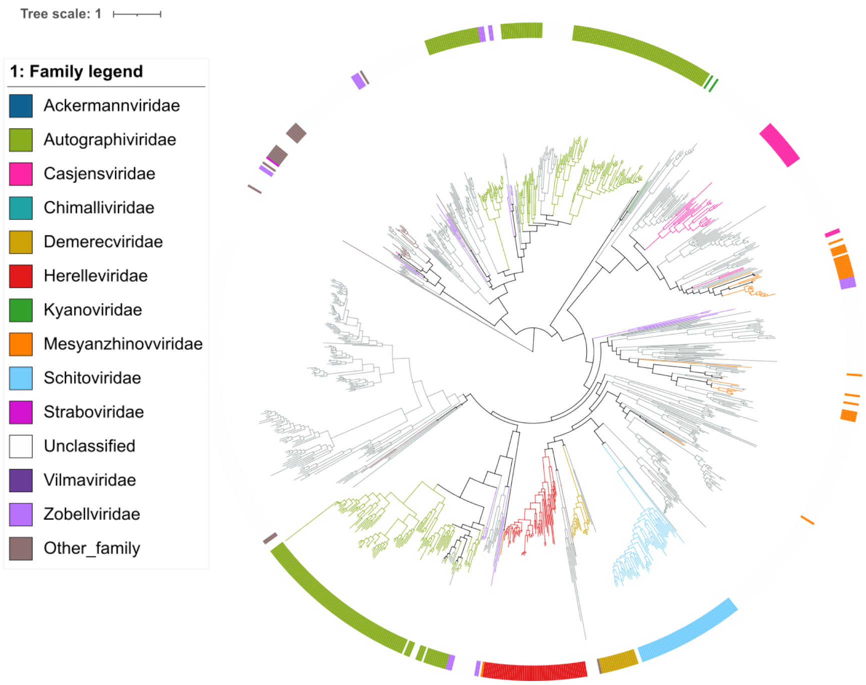


**Supplemental Figure S4.** DNA polymerase A phylogeny. Branches and outer strip colors correspond to family.

### Supplemental Tables

**Supplemental Table S1.** Number of VOGs found in a given percent of Caudo genomes and tree quality values from phylogenetic reconstruction with inCaudo genomes.

| Percent of genomes | Number of VOGs | Abbrev | Tree Certainty | Relative Tree Certainty (0-1) |
|---|---|---|---|---|
| 0.25 | 4042 | 025p | 1621.83298 | 0.641548 |
| 0.5 | 2114 | 05p | 1630.17211 | 0.644847 |
| 1 | 1032 | 1p | 1545.86242 | 0.614902 |
| 2 | 472 | 2p | 1292.08113 | 0.513138 |
